# The entomopathogenic nematode *Steinernema hermaphroditum* is a self-fertilizing hermaphrodite and a genetically tractable system for the study of parasitic and mutualistic symbiosis

**DOI:** 10.1101/2021.08.26.457822

**Authors:** Mengyi Cao, Hillel T. Schwartz, Chieh-Hsiang Tan, Paul W. Sternberg

## Abstract

Entomopathogenic nematodes, including *Heterorhabditis* and *Steinernema*, are parasitic to insects and contain mutualistically symbiotic bacteria in their intestines (*Photorhabdus* and *Xenorhabdus,* respectively) and therefore offer opportunities to study both mutualistic and parasitic symbiosis. The establishment of genetic tools in entomopathogenic nematodes has been impeded by limited genetic tractability, inconsistent growth *in vitro*, variable cryopreservation, and low mating efficiency. We obtained the recently described *Steinernema hermaphroditum* strain CS34 and optimized its *in vitro* growth, with a rapid generation time on a lawn of its native symbiotic bacteria *Xenorhabdus griffiniae*. We developed a simple and efficient cryopreservation method. Previously, *S. hermaphroditum* isolated from insect hosts was described as first-generation hermaphroditic and second-generation gonochoristic. We discovered that CS34, when grown *in vitro,* produced consecutive generations of autonomously reproducing hermaphrodites accompanied by rare males. We performed mutagenesis screens in *S. hermaphroditum* that produced mutant lines with visible and heritable phenotypes. Genetic analysis of the mutants demonstrated that this species reproduces by self-fertilization rather than parthenogenesis and that its sex is determined chromosomally. Genetic mapping has thus far identified markers on the X chromosome and three of four autosomes. We report that *S. hermaphroditum* CS34 is the first consistently hermaphroditic entomopathogenic nematode and is suitable for genetic model development to study naturally occurring mutualistic symbiosis and insect parasitism.

## Introduction

Microbial symbiosis ranges across a wide spectrum from mutualistic (beneficial) interactions to parasitism (Fig. 1A). Mutualistic and parasitic relationships are ubiquitous on earth in all animals and are crucial to the metabolism, immune responses, development, behaviors, which ultimately impact the health and shape the evolution of host animals (Ruby, 2008; McFall-Ngai *et al*., 2013; Morais *et al*., 2021). Currently, studies of the genetic basis for multicellular partners engaging in mutualistic interactions are mostly restricted to traditional model organisms with simplified consortia or synthetic microbiomes, such as hydra, fruit fly, zebrafish, mice, and more recently microbiome studies of the free-living soil nematode *Caenorhabditis elegans* (Bosch *et al*., 2019). Genetic tools have not existed with which to systematically study the animal host side of any naturally-occurring interactions with their mutualistic microbes, especially in a binary and species-specific manner.

**Figure 1:**
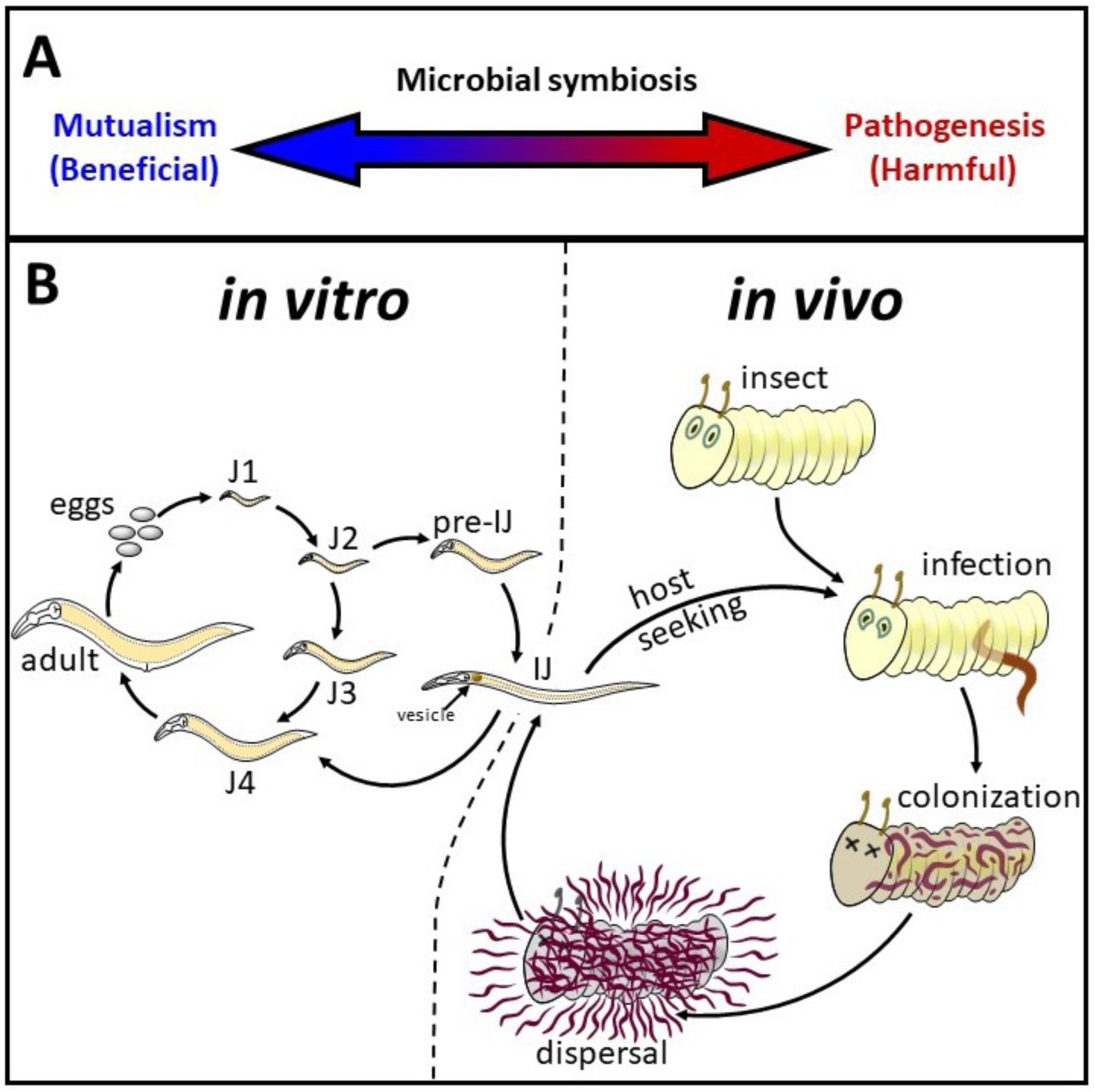
Mutualistic and parasitic life cycle of *Steinernema hermaphroditum*. (A) The relationship between a microbe and its animal host can be considered to be on a continuum between mutualism, from which both organisms benefit, and pathogenesis, in which one species is entirely harmful to the other. (B) The life cycle of the entomopathogenic nematode *S. hermaphroditum*. The reproductive development of *S. hermaphroditum* can be monitored during growth *in vitro*, on Petri plates: the worm passes through four larval stages, J1 to J4, then becomes an adult, which produces fertilized embryos in eggsthat can be laid or can hatch internally, to begin the cycle again. The second larval stage can alternatively develop into a developmentally arrested, stress-resistant infective juvenile (IJ), analogous to the dauer stage of *C. elegans*. The infective juvenile has a specialized vesicle in its anterior intestine containing viable cells of its bacterial symbiont *X. griffiniae*. The IJ will seek out an insect host, invade its body, and resume development, re-entering the reproductive life cycle. As it exits developmental arrest the worm will release its bacterial cargo. Together the worm and its bacteria kill the insect, and the bacteria convert the carcass into a nutritive food source for the worm and for generations of its progeny. When the food source is exhausted a new generation of worms will develop as IJs and will seek out new hosts.

*C. elegans* has been a highly productive laboratory organism for investigating the genetic bases underlying a broad range of biological questions including how genes specify complex structures and behaviors (Brenner, 1974). An equally tractable nematode system with a naturally occurring species-specific microbial symbiosis could be similarly productive in studying symbiotic interactions. The reproductive mode of *C. elegans* consists of self-fertilizing hermaphrodites and males, greatly facilitating the genetic screens that rapidly produce a large number of mutants (Brenner, 1974). These mutatants provided genetic markers to establish the basic genetic features of the organism and led to the discovery of novel phenotypes and important biological mechanisms (Brenner, 1974; Horvitz, 2003; Fire, 2007; Mello, 2007). Large collections of publicly available mutants and rich genetic tools in the wild-type N2 isolate and diverse natural isolates of *C. elegans* have more recently facilitated the study of the genetics of host animals among their interactions with intestinal microbiomes and bacterial pathogens they encounter in the wild (Frézal and Félix, 2015; Shapira, 2017; Kim and Flavell, 2020; Zhang *et al*., 2021). However, *C. elegans* lacks a stable and species-specific mutualistic symbiont and is free-living rather than a parasite, and so cannot be used to investigate the complete range of symbiotic interactions.

Following the establishment of *C. elegans* as a genetic model system, genetic tools have been developed in other nematode species to study their molecular ecology, evolution, and multi-organism interactions, including predator-prey, necromenic, and parasitic relationships (Kroetz *et al*., 2012; Lightfoot *et al*., 2019; Gang *et al*., 2020; Liu *et al*., 2020). However, the limited number and diversity of nematode species that have been developed for genetic research does not match the potential that nematodes have to increase our understanding of diverse biological fields, including symbiotic and parasitic interactions. For instance, although parasitic nematodes infect all animal species and cause major medical problems, the characterization of molecular mechanisms in nematode infections are restricted to a few species of human parasite due to limitations of genetic tools, ethical concerns, and difficulties of culturing parasitic nematodes *in vitro* (Lok, 2019). Despite the limitations imposed by its free-living, non-parasitic lifestyle in the wild, *C. elegans* is still the major genetic model nematode for most molecular ecology studies. Establishing genetic systems in parasitic nematodes that are culturable and tractable in the laboratory will greatly expand our knowledgeof parasitism and help to identify conserved molecular pathways in the process of infection, such as host-seeking behaviors and parasite-derived immunomodulatory effectors (Gang and Hallem, 2016; Bobardt *et al*., 2020).

*Heterorhabditis* and *Steinernema* entomopathogenic nematodes (EPNs) each have a species-specific association with mutualistic bacteria (*Photorhabdus* and *Xenorhabdus,* respectively; Fig. 1) (Forst and Clarke, 2002; Goodrich-Blair and Clarke, 2007). The symbiotic pairs are used commercially in agriculture as an organic pest control mechanism (Ehlers, 2001; Tarasco *et al*., 2017). The developmentally arrested, stress-resistant infective juvenile (IJ) stage of EPNs, an analog of the dauer stage of free-living nematodes, carry symbiotic bacteria in their intestine as they seek their insect prey. IJs invade an insect and exit from developmental arrest, molting into the fourth juvenile stage (J4) while releasing their symbiotic bacteria into the insect (Ciche and Ensign, 2003; Snyder *et al*., 2007). When delivered internally even a very small number of *Xenorhabdus* and *Photorhabdus* bacteria, on the order of <5 colony-forming units (CFUs), can rapidly kill an insect and reproduce in its cadaver, providing a rich food source for their nematode partner, while inhibiting other microbes and animals from consuming the cadaver (Goodrich-Blair and Clarke, 2007). EPN reproductive development includes four stages of juveniles (J1 to J4, analogous to the four larval stages L1 to L4 of *C. elegans*). Exhaustion of the bacterially colonized insect cadaver as a food source signals the nematodes to cease reproductive development and instead produce IJs after the first juvenile stage (developing from J2 to pre-IJ to IJ; similar to the development of *C. elegans* from L2 to L2d to dauer) and disperse from the insect cadaver seeking new insect hosts (Forst and Clarke, 2002).

The ability to culture EPNs in the laboratory and the availability of genetic manipulation of symbiotic bacteria would make them valuable systems to study fundamental principles of microbial symbiosis ranging from mutualism to parasitism. This includes bacteria-nematode recognition and nutritional codependence, nematode toxin secretion, host-seeking behaviors, and comparative genomics (Herbert and Goodrich-Blair, 2007; Hallem *et al*., 2011; Dillman *et al*., 2012, 2015; Chaston *et al*., 2013; Chang *et al*., 2019). However, no stable genetic tools or comprehensive genetic system has been established in any EPN species despite efforts from multiple groups over decades (Hashmi *et al*., 1995; Ciche and Sternberg, 2007; Ratnappan *et al*., 2016). The lack of efficient cryopreservation methods in EPNs has led to potential issues of inbreeding depression in laboratory strains (Hopper *et al*., 2003). In addition, most of the currently available nematode genetic models took advantage of consistent hermaphroditism similar to *C. elegans,* greatly facilitating screens for homozygous F_2_ progeny of mutagenized animals produced by self-reproduction and the maintenance of mutant lines that might be incapable of mating (Brenner, 1974; Sommer, 2006; Shinya *et al*., 2014). Both *Heterorhabditis* and *Steinernema* lack consistently hermaphroditic species, which has impeded development of forward genetic tools. *Heterorhabditis* alternates between hermaphrodite and female in insects, and when grown on Petri plates produces almost exclusively females after the first generation that recovers from IJ as hermaphrodites, with a low percentage of males that diminishes in successive generations ((Dix *et al*., 1992)HTS and PWS, unpublished). Existing strains of *Steinernema* have been male-female (gonochoristic). The only exception is *S. hermaphroditum-*Indonesia, described as alternating between generations that develop as hermaphrodites and generations that develop as females, similar to *Heterorhabditis* (Stock *et al*., 2004). However this isolate was subsequently lost. To date, forward genetics screens in *Heterorhabditis* have produced only few mutants, insufficient to establish the species as a genetic model (Cohen-Nissan *et al*., 1992; Rahimi *et al*., 1993) HTS and PWS, unpublished), and no attempts to perform forward genetic screens in *Steinernema* have been reported.

In this research, we obtained a recently described isolate of *S. hermphroditum*, the Indian strain CS34 (Bhat *et al*., 2019), and developed protocols for continuous cultivation *in vitro* on agar media in Petri plates. *S. hermaphroditum* can be grown on various bacterial species, including its native symbiont. *In vitro* growth led us to discover CS34 is consistently hermaphroditic, with healthy autonomously reproducing hermaphrodites and spontaneous functional males. We stabilized the genetic background of the wild isolate by repeated inbreeding for controlled number of generations and established efficient cryopreservation to facilitate maintenance of wild-type and collections of mutant strains. We performed ethyl methane sulphonate (EMS) mutagenesis screen and isolated multiple recessive mutants with classical phenotypes such as dumpy (Dpy) and uncoordinated (Unc). Mutant hermaphrodites were mated with wild-type males and heterozygous F_1_ animals produced F_2_ progeny that showed segregation according to Mendelian ratios, proof that these animals reproduce by self-fertilization rather than parthenogenesis. DAPI staining of oocytes of hermaphrodites revealed five pairs of chromosomes. Our initial mapping efforts assigned some of the current mutant alleles into at least four of the five linkage groups, including an X chromosome that determines the sex of this species. Here we report the first consistently hermaphroditic entomopathogenic nematode species, the *S. hermaphroditum* isolate CS34, and demonstrate that it is convenient to culture and cryopreserve in the laboratory and is highly genetically tractable. We propose this species has great potential to further expand our understanding of the genetic bases of a variety of molecular signaling pathways, especially in microbial symbiosis.

## Materials and Methods

### Nematode maintenance and bacterial culture

Conventional (in contrast to germ-free or axenic) infective juveniles are defined as those colonized by their bacterial symbiont. Conventional IJs of *Steinernema hermaphroditum* were propagated through infection of 5^th^ instar larvae of the wax moth *Galleria mellonella* (PetSmart, Phoenix, AZ) and recovered using a modified White trap (White, 1927). IJs were trapped in distilled water and maintained in 15 mL culture flasks (Corning, Corning, NY). To grow *S. hermaphroditum in vitro,* nematodes were grown on bacterial lawns on the surface of agar media in Petri plates. To grow symbiotic bacterial lawns, individual colonies of *X. griffiniae* were picked from Luria Bertani agar supplemented with 0.1% sodium pyruvate (LB-pyruvate) (Xu and Hurlbert, 1990) and used for inoculating liquid media. Cultures were grown in LB broth kept in the dark (dark LB) overnight at 30 °C with shaking or in 2% proteose peptone #3 with 0.1% sodium pyruvate overnight at room temperature without shaking. Bacterial cultures were seeded onto lipid agar media (per liter: 15 g bacto agar, 8 g nutrient broth, 5 g yeast extract, 2 g McCl_2_·6H_2_O, 1 g sodium pyruvate, 7 ml corn syrup, 4 ml corn oil (Vivas and Goodrich-Blair, 2001) or nematode growth media (NGM) agar (Brenner, 1974).

### Symbiotic bacteria isolation and nematode growth

*S. hermaphroditum* IJs were concentrated by centrifugation, surface-sterilized by treatment with 1% bleach for two minutes with gentle shaking, concentrated by centrifugation, and washed in distilled water three times. Approximately 200 IJs were resuspended in 200 µL dark LB media and ground for two minutes using a hand homogenizer (Kimble Pellet Pestle Cordless Motor, DWK Life Sciences, Millville, NJ). Samples were examined using microscopy to confirm that grinding of all IJs was complete. Homogenate was serially diluted to 1:10, 1:100, 1:1,000, and 1:10,000 in dark LB. Each dilution was plated on LB-pyruvate agar and incubated at 30°C overnight. Dilutions that produced 10-100 colonies were counted to estimate colony forming units (CFU) per IJ. Ten individual colonies were purified by re-streaking onto LB-pyruvate agar followed by isolating single colonies. Symbiotic bacterial isolates were frozen at - 80°C after mixing 900 µL of overnight bacterial culture with 600 µL of LB-glycerol (50% LB, 50%glycerol). One *X. griffiniae* isolate was given the name HGB2511 (Table S1) and was used as the principal food source for culturing *S. hermaphroditum* on agar media.

### 16S rRNA gene sequence determination from symbiotic bacterial isolates

To extract genomic DNA, overnight cultures of symbiotic bacteria were pelleted by centrifugation at maximum speed in a table top microcentrifuge (∼16,000 rcf) for 1 minute. The bacterial pellet was resuspended in 50 µL distilled water, boiled at 100°C for 10 minutes, incubated on ice for 10 minutes, and centrifuged at maximum speed in a microcentrifuge remove debris. Supernatant containing genomic DNA was used as a template for the PCR amplification of the 16S rRNA gene using *Taq* polymerase (New England Biolabs, Ipswich, MA) and the primers oMC98f (5’-GAAGAGTTTGATCATGGCTC-3’) and oMC99r (5’-AAGGAGGTGATCCAGCCGCA-3’) as previously described (Tailliez *et al*., 2006). An approximately 1500bp PCR product was isolated using agarose gel electrophoresis and purified using QIAprep (QIAGEN, Valencia, CA). Sanger-sequencing was performed by Laragen (Culver City, CA) using primers oMC98f, oMC99r, oMC100r (5’-ACCGCGGCTGCTGGCACG-3’), and oMC101r (5’-CTCGTTGCGGGACTTAAC-3’). The 16S rRNA gene sequence was compared to existing rRNA gene sequences in the NCBI database using BlastN (NCBI Resource Coordinators, 2017).

### Growth rate analysis under different culture conditions

Embryos of *S. hermaphroditum* were obtained by treating gravid hermaphrodites with NaOH and NaOCl similar to methods used for *C. elegans* (Stiernagle, 2006). The centrifugation process of embryo extraction was performed using a clinical centrifuge with a uniform speed. Briefly, gravid hermaphrodites cultured with *X. griffiniae* on NGM at 25°C were washed off the plates with M9 buffer (per liter: 3 g KH_2_PO_4_, 6 g Na_2_HPO_4_, 5 g NaCl, with 1 ml 1 M MgSO_4_ added after autoclaving). The nematode suspension was then centrifuged to remove the supernatant, and water was added for a total volume of 3.5 mL. 0.5 mL of 5M NaOH and 1 mL of household bleach (8% available chlorine) were then added to the solution. The solution was mixed by gently shaking and allowed to react for 4-6 minutes, after which the embryos were collected by centrifugation. Embryos were then washed with 10 mL of M9 buffer three times, then centrifuged to remove the supernatant. Embryos were resuspended in M9 and seeded onto NGM agar Petri plates on which bacterial lawns had been grown. For growth rate analysis of worms fed with different bacteria, the bacterial strains used were *Xenorhabdus griffiniae* HGB2511, *Comamonas aquatica* DA1877, *Escherichia coli* OP50, and *Escherichia coli* HB101 (Table S1). Petri plates containing the bacterial lawns and embryos were cultured at 25°C. To assess the effects of temperature on growth, embryos were placed on *X. griffiniae* HGB2511 bacterial lawns and incubated at 20°C, 25°C, 27.5°C, and 30°C. Nematodes on these plates were imaged every 12 hours using WormLab equipment and software (MBF Bioscience, Williston, VT) until 120 hours had elapsed unless the animals were first obscured by the growth of their progeny. Body length was measured by drawing a line from head to tail through the midline of each worm using ImageJ (NIH) software similar to an established method (Roh *et al*., 2012).

### Cryopreservation of *S. hermaphroditum*

*S. hermaphroditum* cryopreservation was adapted from a trehalose-DMSO freezing protocol for *C. elegans* (O’Connell, 2021). Briefly, nematodes were grown on NGM agar seeded with *X. griffiniae* bacteria until the food was nearly exhausted. A mixed stage population of juvenile nematodes (mostly J1 nematodes that had hatched internally inside their hermaphrodite mothers) were washed off the agar in M9 buffer. Nematodes were concentrated by centrifugation for 1 minute at approximately 1,000 rcf and washed in 5 mL of trehalose-DMSO solution (per liter: 30.2 g trehalose, 35.4 mL DMSO, filter-sterilized), then incubated in 5 mL trehalose-DMSO solution for more than 30 minutes. Samples were transferred into cryotubes, placed in sealed Styrofoam boxes to slow their temperature change, and then placed in a -80°C freezer overnight before being transferred to cardboard boxes in a -80°C freezer.

### Dissection and staining of the *S. hermaphroditum* gonad

A protocol for dissection and staining of the *S. hermaphroditum* gonad was adapted from gonad dissection and staining protocols developed for *C. elegans* (Kocsisova *et al*., 2018, 2019). Briefly, young adult hermaphrodites (one day post J4) were picked into a watchglass (Carolina Biological Supply Company, Burlington, NC) containing phosphate-buffered saline (PBS). Worms were washed three times on the watchglass with PBS and immobilized with levamisole (final concentration 200 µM). The worms were then dissected at the pharynx with a pair of 30G 5/8” needles (PrecisionGlide, BD, Franklin Lakes, NJ). The dissected gonads were fixed with 3% paraformaldehyde (EM Grade, Electron Microscopy Science, Hatfield, PA) in PBS and post-fixed with 100% methanol. Fixed gonads were then rehydrated and washed three times with PBSTw (PBS + 0.1% Tween-20). Following the washes, the gonads were incubated overnight at room temperature with a monoclonal anti-MSP antibody (4A5, Developmental Studies Hybridoma Bank, University of Iowa; Kosinski *et al*., 2005) diluted 1:10 in 30% goat serum (Thermo Fisher Scientific, Waltham, MA) in PBS. The gonads were then washed three times with PBSTw and incubated with secondary antibodies (Goat anti-Mouse Alexa Fluor 555; A-21424, Invitrogen) diluted 1:400 in 30% goat serum in PBS for 4 hours at room temperature. Following the secondary staining, the gonads were again washed three times with PBSTw and resuspended in 1 drop of Vectashield containing 4’,6-diamidino-2-phenylindole (DAPI) (Vector Laboratories, Burlingame, CA). The gonad was finally mounted on pads of 5% agarose in water on microscope slides and covered with a cover glass (Sulston and Horvitz, 1977; Kocsisova *et al*., 2018). For gonads with only DAPI staining, antibody staining steps were skipped.

### Image acquisition

Photomicrographs in Figure 2B, 2C and Figure 4 were acquired with a Zeiss Imager Z2 microscope equipped with Apotome 2 and Axiocam 506 mono using Zen 2 Blue software. Figure 3A-3D were acquired through the WormLab (MBF Bioscience, Williston, VT) equipment and software. The camera was a Nikon AF Micro 60/2.8D with zoom magnification.

**Figure 2:**
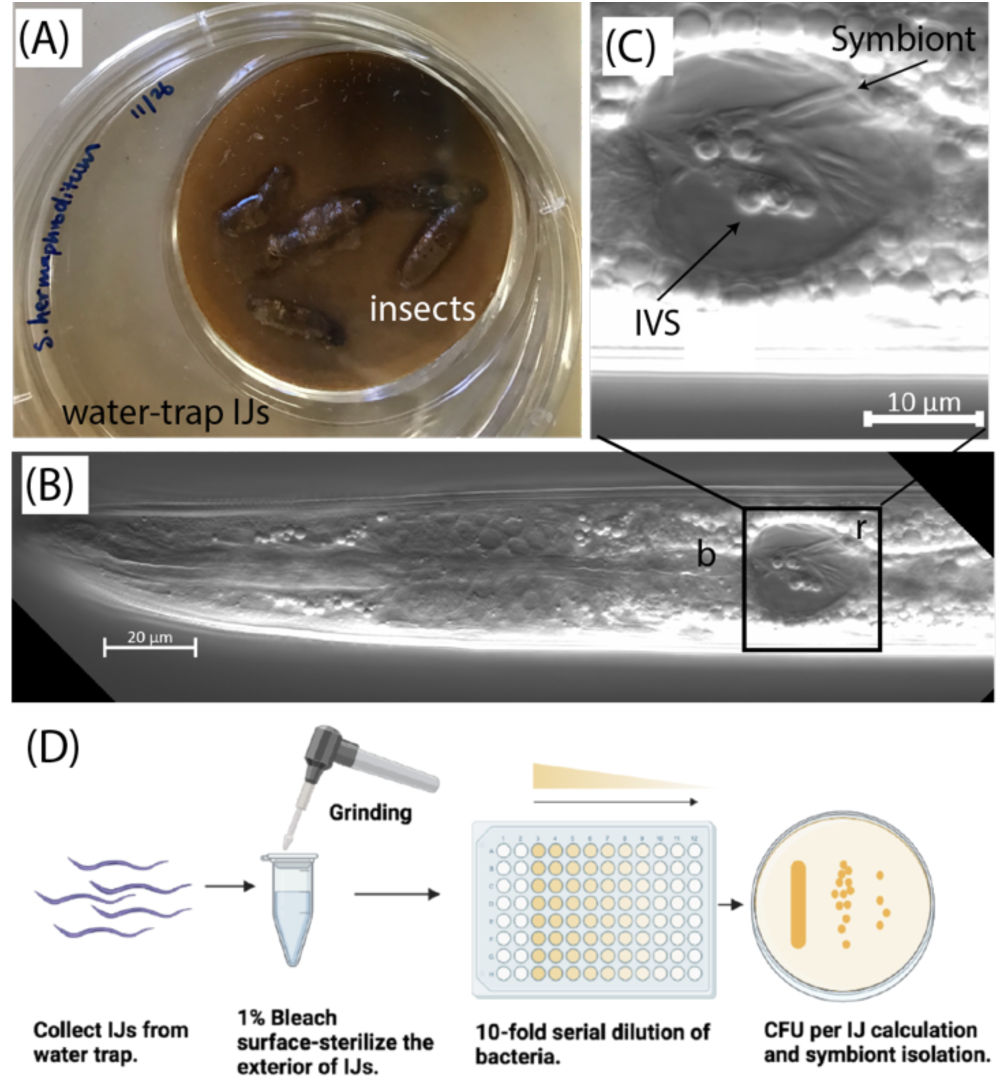
*In vivo* growth of *S. hermaphroditum* and symbiotic bacteria isolation. (A): A modified White trap for the recovery of infective juvenile (IJ) stage larvae as they disperse from colonized carcasses of *Galleria mellonella* wax moth larvae. (B): Photomicrograph using Nomarski differential interference contrast optics to show the head of an IJ stage *S. hermaphroditum*. Box labeled “r” indicates the receptacle, an intestinal pocket colonized with native symbiotic bacteria. “b” indicates the position of the basal (posterior) bulb of the pharynx. Scale bar, 20 um. (C): An expanded view of the intestinal vesicle in panel B. Rod-shaped symbiotic bacteria are localized in the lumen of the intestinal vesicle. Some bacterial cells adhere to a spherically shaped intravesicular structure (IVS). Scale bar, 10 μm. (D): Isolation and quantitfication of symbiotic bacteria that have colonized the intestines of *S. hermaphroditum* IJ larvae. IJs are collected from infected insect hosts, treated with bleach to kill any bacteria on their surfaces, and ground to release bacteria within their intestines. These bacteria are serially diluted until individual colonies can be counted and recovered for further analysis.

**Figure 3:**
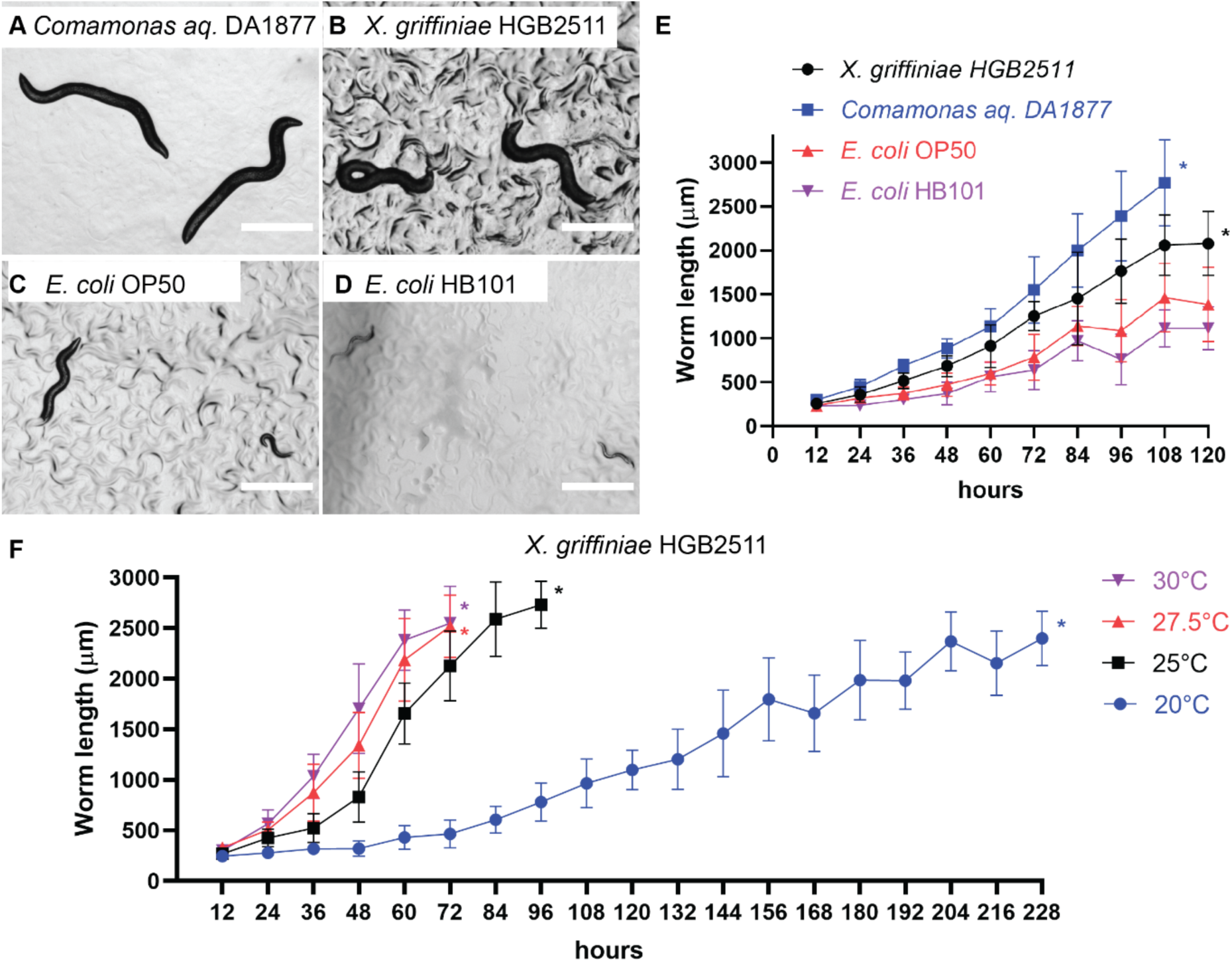
*In vitro* growth of *S. hermaphroditum*. (A-D). *S. hermaphroditum* embryos free of bacteria were obtained by dissolving gravid adults in bleach solution. The embryos were then seeded on NGM agar plates with lawns of different bacteria as a food source. Images show the sizes of worms that have been grown from embryos at 25°C for 96 hours. Scale bars indicate 1 mm. Bacterial lawns were: (A) The *S. hermaphroditum* CS34 native symbiont. *Xenorhabdus griffiniae* HGB2511; (B) *Comamonas aquatica* DA1877; (C), *E. coli* OP50; (D) *E. coli* HB101.(E-F). Quantification of nematode growth. Body length was measured along the midline of the animal as described in Materials and Methods. Asterisks indicate the time point after which the experiment was terminated because animals were obscured by their progeny. (E) Quantification of *S. hermaphroditum* growth on different bacterial lawns on NGM at 25°C. Worms grown on *C. aquatica* were significantly larger than those grown on *X. griffiniae* at every time point (p < 0.01, Student’s t-test), while worms grown on the two *E. coli* lawns were smaller than those grown on *X. griffiniae*, a difference that was statistically significant at every time point except for OP50 at 24 hours (p < 0.05, Student’s t-test). Values are mean ± SD (n = 17 - 20) (F) Quantification of *S. hermaphroditum* growth on its native symbiont *X. griffiniae* HGB2511 at different temperatures. *S. hermaphroditum* growth is strongly influenced by temperature. Worms grown at 27.5°C and 30°C were significantly (p < 0.05, Student’s t-test) larger than those grown at 25°C at every time point assayed, while worms grown at 20°C were significantly smaller (p<0.01, Student’s t-test). Values are mean ± SD (n = 20)

**Figure 4:**
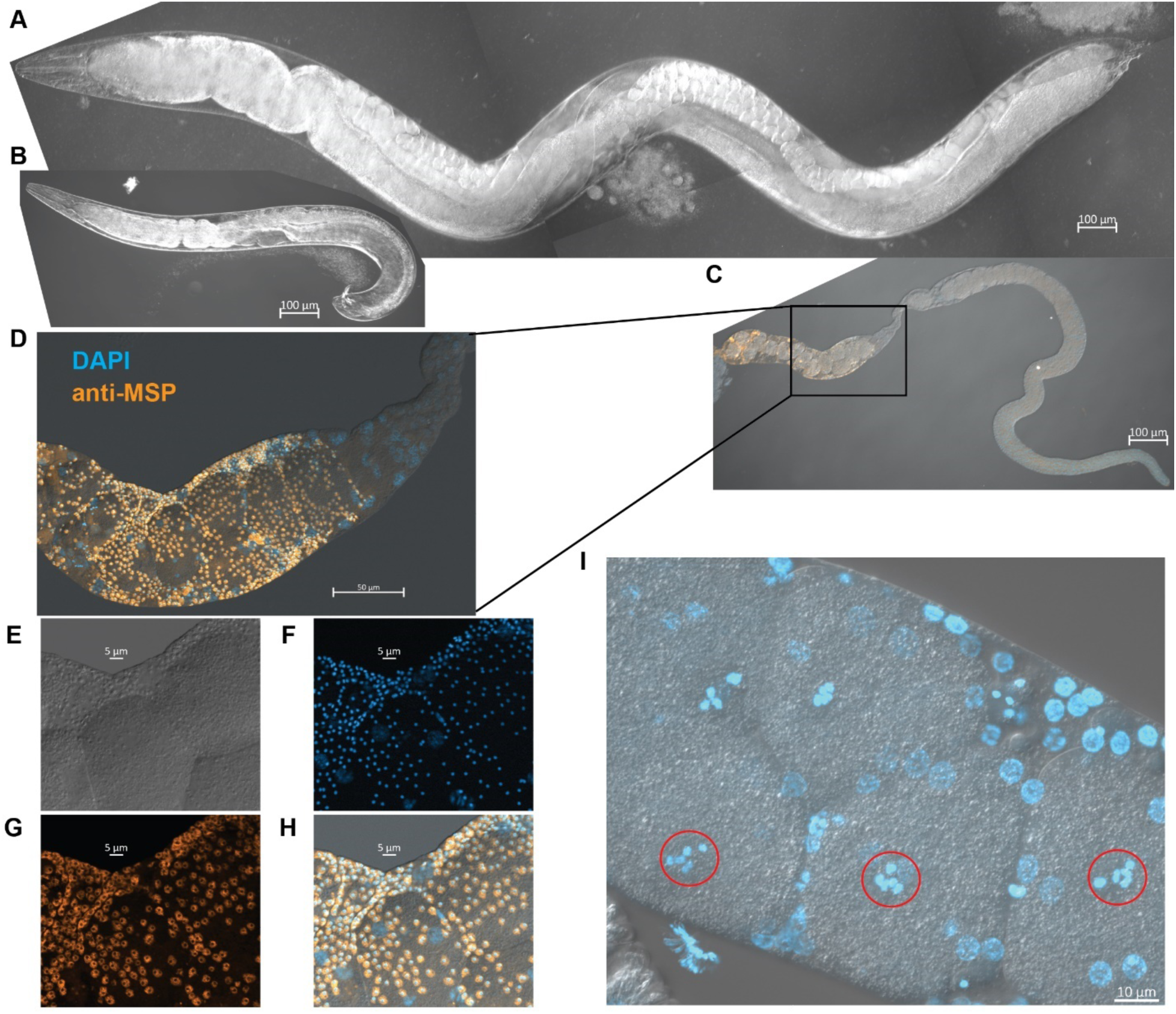
*S. hermaphroditum-*India is consistently hermaphroditic. Progeny of an unmated *S. hermaphroditum* hermaphrodite consist mostly of hermaphrodites but also include rare males. **(A)** An adult hermaphrodite of *S. hermaphroditum.* Scale bar, 100 μm. **(B)** An adult male of *S. hermaphroditum* at the same magnification. Scale bar, 100 μm. **(C)** A dissected gonad arm from a young hermaphrodite stained with the DNA dye DAPI (blue) and with an antibody against Major Sperm Protein (MSP) (orange). Scale bar, 100 μm. **(D)** A section of the gonad arm shown in (C) with higher magnification showing the presence of sperm (orange) in the spermathecal of an unmated hermaphrodite. Scale bar, 50 μm. **(E-H)** Higher magnification images of a section of the spermatheca showed in (D) The sperm of *S. hermaphroditum* appear to be small, especially as seen using DIC. (E) Nomarski microscopy (differential interference contrast, DIC); (F) DAPI; (G) Anti-MSP; (H) Merged. Scale bars, 5 μm. **(I)** Oocytes in diakinesis show five pairs of tetraploid chromosomes at Meiosis Prophase I (red circles). Scale bar, 10 μm.

### EMS mutagenesis and screening for visible mutants

Ethyl methanesulfonate (EMS) mutagenesis of *S. hermaphroditum* was adapted from the standard protocol used for *C. elegans* (Brenner, 1974). Unseeded and *X. griffiniae* seeded NGM agar were used throughout the EMS screen, and will be referred to as unseeded plates and bacterial lawn, respectively. Animals were hand-picked by developmental stage onto an unseeded plate, suspended in M9 buffer, and collected in a 15 mL conical tube. Nematodes were rinsed with M9 and incubated for four hours at 20°C in 4 mL of M9 containing 20 μL EMS (46 mM), rotating on a cell culture wheel to avoid settling. Animals were pelleted by centrifugation (1 minute, approximately 1000 rcf) and rinsed several times with M9, then allowed to recover at least 30 minutes at room temperature on bacterial lawn. After recovery, animals were transferred to new bacterial lawns with one to five animals per plate. Mutagenesis was performed using mid-J4 larvae, late-J4 larvae, and young adult hermaphrodites in our initial trials. Young adults were found to be the most productive stage to mutagenize.

The F_2_ progeny of mutagenized animals were examined for possible mutant phenotypes (Fig. 5A), either on the same plate the mutagenized animals had been growing on, or by transferring the progeny of the mutagenized animals to new plates to avoid exhausting the food source. When transferring, animals were either washed off the plate using M9 or moved within a chunk of agar cut by heat-sterilized metal spatula. Nematodes were then placed on the surface of another larger Petri plate (10 cm in diameter) containing NGM agar seeded with *X. griffiniae*. Mutants were tracked to identify the mutagenized P_0_ animal or animals from which they descended, to avoid repeated recovery of animals with the same phenotype that might have arisen from the same mutagenesis event. Candidate mutants were picked to bacterial lawn and their self-progeny were examined for the propagation of a mutant phenotype. Stable mutant lines were mated with wild-type males to test for recessiveness and for linkage to the X chromosome.

**Figure 5:**
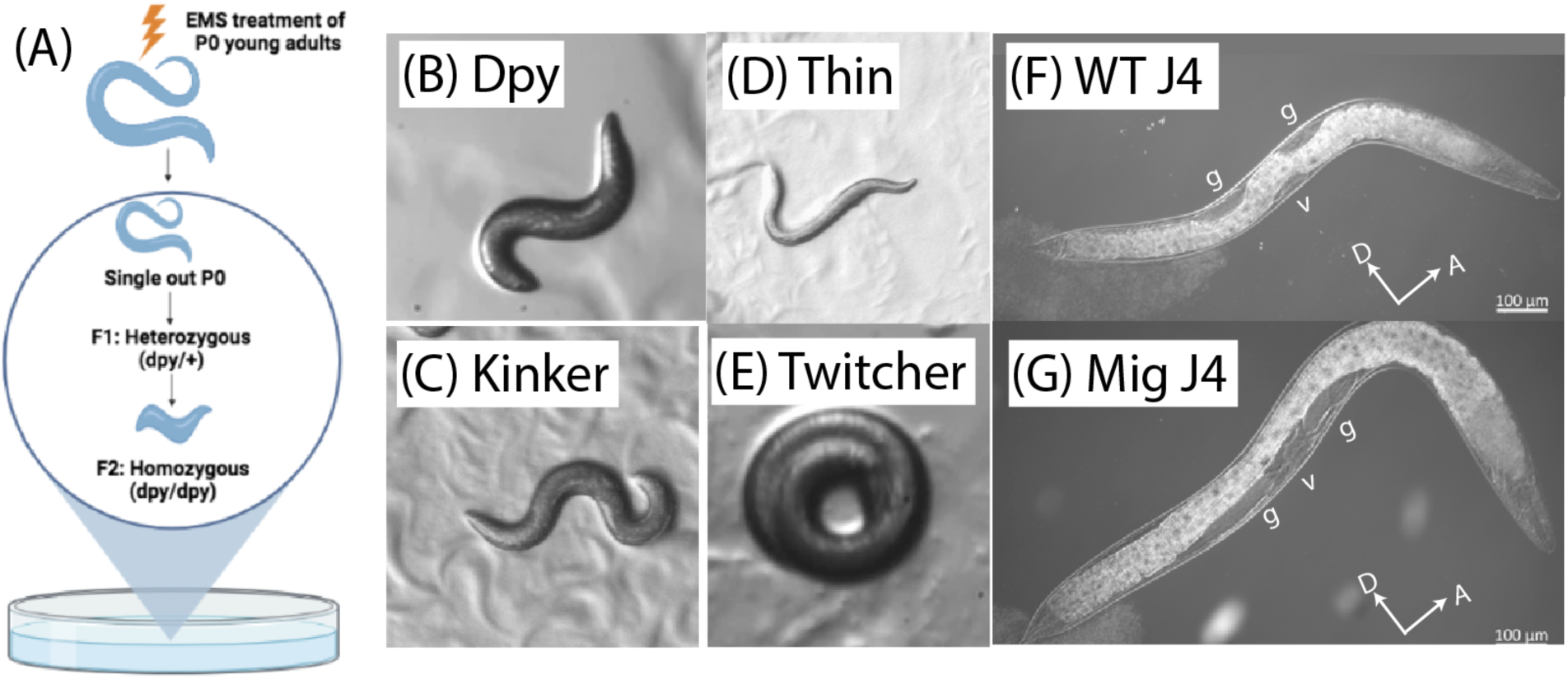
EMS mutagenesis of *Steinernema hermaphroditum.* (A): EMS mutagenesis of young adults (the P_0_ or parental generation) and isolation of mutant lines from individuals in the F_2_ generation. Candidate mutant strains are tracked to identify the mutagenized P_0_ animal they descended from. (B-G): examples of mutant phenotypes. (B): The dumpy appearance of *dpy(sy1646).* (C): The resting posture of the uncoordinated mutant *unc(sy1636)*, a kinker Unc. (D): The long, thin uncoordinated mutant *unc(sy1653).* (E): The twitcher mutant *unc*(*sy1654*); see also a film of its phenotype in supplementary online materials. (F): Wild-type (WT) fourth-stage J4 larva. The developing gonad starts on the ventral side near the animals midpoint (labeled “v”) and expands both anteriorly and posteriorally. As the gonad expands it pushes aside the intestine (which has a white appearance), resulting in clear patches (labeled “g”). It can be seen that the gonad has moved dorsally both anterior and posterior to its starting point. Anterior is to the right and dorsal is up. Scale bar, 100 μm. (G): Altered gonad migration is apparent in the J4 stage of a *mig(sy1637)* mutant. Note that the expanding gonad has remained entirely ventral as it has expanded to the anterior and the posterior of its starting point (labeled “V”). Anterior is to the right and dorsal is up. Scale bar, 100 μm. “A” denotes for anterior; “D” denotes for dorsal; “v” indicates the position of vulva; and “g” indicates the position of gonad arms.

To test for linkage between two mutations, we generated animals carrying both mutations as heterozygotes with wild-type alleles, in *trans* to each other if they were linked. Except in rare cases of mutants that could mate as homozygous or hemizygous males, this was done by first mating wild-type males to one of the two mutant strains. We then recovered F_1_ males from the cross and mated them to the second mutant strain. Individual cross progeny from this second cross were placed on NGM agar Petri plates seeded with *X. griffiniae* and allowed to propagate. All of these animals would be heterozygous for the second mutation, but only half would carrying the first mutation; these animals were identified by examining their progeny. From plates derived from animals heterozygous for both mutations we identified animals whose phenotype indicated they were likely to be homozygous for one of the two mutations present and placed each onto a bacterial lawn. By examining the progeny of these animals we first confirmed that they were homozygous for the mutation selected and then scored for animals phenotypic for the other mutation, which would mean their parent had one parental genotype and one recombinant genotype. Unlinked mutations would be expected to be present 2/3 of the time; tightly linked mutations should rarely be present. This approach was chosen because not all mutant phenotypes could be unambiguously scored in individual animals, and homozygosity for one mutation would often make it difficult to score for homozygosity of the other.

### Statistics

Animal sizes were compared using the two-tailed unpaired Student’s t-test. All animal sizes are mean ± SD, n = 17 - 20

## Results

### Symbiotic bacteria isolation and *In vitro* growth of *S. hermaphroditum*

Consistent *in vitro* growth of nematodes is crucial to optimizing the health, life cycle progression, mating efficiency, and visibility of animals, ultimately facilitating genetic study of the organism (Nigon and Félix, 2017). Unlike some other nematodes that could breed on a standard *E. coli* lawn on NGM agar, *Steinernema* entomopathogenic nematodes usually require their symbiotic bacteria to achieve optimal growth *in vitro* unless they are provided with non-transparent and complex media (Flores-Lara *et al*., 2007; Murfin *et al*., 2012). We collected *S. hermaphroditum* IJs emerged from parasitized *Galleria mellonella* insect larvae and visualized symbiotic bacteria localized in a specialized intestinal pocket termed the receptable (previously described as the vesicle; Fig. 2A and 2B; (Bird and Akhurst, 1983; Kim *et al*., 2012). Contained in the luminal space of the receptacle, some symbiotic bacteria adhere to an intravesicular structure (IVS), an untethered cluster of spherical bodies that has been reported in other *Steinernema spp* (Fig. 2C; (Martens and Goodrich-Blair, 2005; Sugar *et al*., 2011). To isolate native symbiotic bacteria from *S. hermaphroditum* we first used a 1% bleach solution to sterilize the exteriors of IJs and then extracted their symbiotic bacteria by grinding (Fig. 2D). We estimated the average IJ contained 5-10 bacterial colony forming units (CFUs) capable of growing on LB-pyruvate agar. *Xenorhabdus* bacteria can exhibit colony-to-colony phenotypic variation that can affect the reproduction of nematodes feeding on them (Boemare and Akurst, 1988; Volgyi *et al*., 1998; Park *et al*., 2007; Sugar *et al*., 2011; Cao *et al*., 2017). To examine a range of possible phenotypes from the bacterial extraction, we isolated ten individual colonies as candidate strains for feeding nematodes *in vitro.* Eight isolates produced brown pigment on NGM agar while the other two isolates did not. Similar phenotypic differences in pigment production were reported in other *Xenorhabdus spp* as a signature of phenotypic variation (Sugar *et al*., 2011). To identify the symbiont isolates we amplified 16S rRNA gene and determined that all ten isolates had the same sequence, one that is more than 99% identical to the sequence previously reported for *Xenorhabdus griffiniae* strain ID10^T^, a symbiotic bacterial strain isolated from a subsequently lost Indonesian isolate of *S. hermaphroditum* (Stock *et al*., 2004; Tailliez *et al*., 2006). Therefore, the phenotypic differences among bacterial isolates is not caused by the co-existence of multiple species of symbiont, but more likely due to phenotypic switching within the same strain, or multiple strains of *X. griffiniae* co-existing in the same natural population of nematode.

To culture *S. hermaphroditum* nematodes *in vitro,* we first compared the growth of these nematodes on their native symbiotic bacteria on three growth media: lipid agar, traditionally used in EPN growth (Vivas and Goodrich-Blair, 2001); NGM agar, widely adopted for *Caenorhabditis* and other free-living nematodes (Brenner, 1974); and NGM agar supplemented with 0.1% pyruvate, which optimizes the growth of some species of *Xenorhabdus* bacteria (Fig. S1) (Xu and Hurlbert, 1990). Symbiotic bacteria grew sufficiently on all three media to support the growth of *S. hermaphroditum.* The nematodes were observed to progress through the stages of their reproductive life cycle (J1-J2-J3-J4-Adult-embryo; see Fig. 1B) at a similar pace on all three media in the first generation of growth, while the nutrient-rich lipid agar supported additional generations of nematode growth that the other two media did not (Fig. S1A). IJs placed on bacterial lawns recovered from developmental arrest and resumed reproductive development on all three media but at slightly different paces (Fig. S1B). We selected nematode growth media (NGM) agar as the standard growth condition for genetics study; it is more transparent than lipid agar and supports a thinner bacterial lawn that permits easier visualization and manipulation of the nematodes within it (Fig. 3A). To expand our repertoire of *in vitro* growth conditions, we further tested nematode growth feeding on bacterial strains commonly used in the *C. elegans* research community, including *Comamonas aq* DA1877 (Fig. 3B; (Shtonda and Avery, 2006; Watson *et al*., 2014), *E. coli* OP50 (Fig. 3C; (Brenner, 1974), and *E. coli* HB101 (Fig. 3D; (Boyer and Roulland-dussoix, 1969). We found the native symbiont *X. griffiniae* better supported nematode growth *in vitro* than did either of the two *E. coli* strains tested (Fig. 3E). *S. hermaphroditum* nematodes grew faster on *Comamonas* than on their native symbiont (Fig. 3E), suggesting that *Comamonas* amply provides nutrients essential for *S. hermaphroditum* development, that are provided by *E. coli* in limited amounts.

In the laboratory, EPN species have usually been cultured at 20-27°C similar to other soil nematodes (Dunphy and Webster, 1989). Our initial trial of breeding *S. hermaphroditum* at room temperature resulted in slow growth (approximately 10 days per generation), while higher temperatures supported higher fertility and faster development, consistent with the tropical environment from which they were isolated (Fig. S1C and S1D). At 33°C the hermaphrodites cannot produce living embryos, while 37°C causes lethality of the animals (Fig. S1C and S1D). To better optimize the growth temperature, we monitored nematode growth under various temperatures from 20-30°C (Fig. 3F). *S. hermaphroditum* CS34 showed a range of generation times: roughly 2.5 days at 30°C, 3 days at 27.5°C, 4 days at 25°C, and more than 9 days at 20°C. The generation time at 25-30°C is similar to the generation time of *C. elegans* growth in the laboratory at its optimal cultivation temperature. We used 25°C as a standard growth condition for *S. hermaphroditum* to facilitate the adaptation of breeding and genetic techniques developed for use in *C. elegans*.

### Establishing a protocol for cryopreservation of *S. hermaphroditum*

The short generation time of nematodes and the lack of outbreeding in a selfing hermaphrodite mean that continuously cultured *S. hermaphroditum* would accumulate mutations and might rapidly show inbreeding depression (Dolgin *et al*., 2007). Cryopreservation is therefore a crucial technique to maintain healthy lines. To develop a cryopreservation method, we first attempted to freeze *S. hermaphroditum* IJs adapting a methanol-wash protocol that has been previously adopted in multiple *Heterorhabditis* and *Steinernema* species (Popiel and Vasquez, 1991) but failed to consistently recover viable nematodes from frozen stocks. An attempt using a glycerol-based freezing protocol adapted from *C. elegans* research gave a low yield of viable *S. hermaphroditum* that survived freezing (5-20 nematodes per mL) (Table S2). We more successfully adapted a recently reported trehalose-DMSO protocol for freezing *C. elegans* research (O’Connell, 2021). Using this protocol we efficiently froze and recovered *S. hermaphroditum* mixed stage juveniles (Table S2). Nematodes grown on NGM showed higher efficiency (>200 nematodes survived per mL) than did those grown on lipid agar (5-20 nematodes per mL), suggesting the two growth conditions may cause drastic physiological differences. The frozen stock was test-thawed over the course of two months and was found to provide stable, highly efficient recovery. We used the resulting trehalose-DMSO freezing protocol for the cryopreservation of *S. hermaphroditum* strains in this research.

### Establishing a homozygous and stable isogenic line via inbreeding

A population of nematodes isolated from the wild is likely to contain a mix of genotypes and to be heterogeneous in genome sequences (Barrière and Félix, 2006). To improve future genome sequence assembly and to provide a genetically and phenotypically consistent population for genetic study, it is useful to obtain a homozygous and stable isogenic line via inbreeding (Barrière and Félix, 2006). *S. hermaphroditum* was previously described to be gonochoristic for alternating generations, which is more likely to cause inbreeding depression when immediately isolating single mating pairs (Stock *et al*., 2004). We therefore adopted an inbreeding strategy appropriate to a male-female population by first establishing ten mating groups of nematodes co-cultured with ten symbiotic bacterial isolates (Ancestral groups I to X corresponding to symbiotic bacteria isolates 1-10, respectively, Fig. S2) (Barrière and Félix, 2006). We discovered that feeding on the eight pigmented (brown) bacterial isolates (Ancestral groups I, II, III, V, VI, VIII, IX, and X) produced relatively higher progeny numbers, while feeding on either of the non-pigmented bacterial isolates (Ancestral groups IV and VII) produced relatively fewer progeny (data not shown). We chose five ancestral lines including both pigmented and non-pigmented symbionts (Ancestral groups III, IV, VI, VII, and IX) for further inbreeding by isolating ten single mating pairs from each of the five ancestral lines. During this process of inbreeding, we observed a low (less than 5%) frequency of males in the population and found that the animals with a grossly female appearance could consistently reproduce when males were absent (see details below). Having found that the *S. hermaphroditum* wild isolate CS34 is stably hermaphroditic for multiple generations, we inbred the wild type line by isolating single self-reproducing virgin juvenile hermaphrodites for five to ten consecutive generations on various lines of symbiotic bacteria (Fig. S2, Table S1). One strain, PS9179, derived from ancestral line IX, inbred by selfing single animals for ten generations co-cultured with the pigmented bacterial isolate HGB2511, was chosen to function as a reference strain. Animals of this strain are healthy and active, and have retained their ability to infect and kill *Galleria mellonella* insect larvae and reproduce in the resulting carcass, eventually producing a new generation of infective juveniles.

### *S. hermaphroditum* is consistently hermaphroditic

Next we examined if sperm was present in unmated animals with female-like somatic morphology, which would be evidence of hermaphroditism. We dissected the gonads of virgin hermaphrodites and observed sperm-like objects in a region of the gonad that corresponds to the spermatheca of other species (Fig. 4C and 4D). The sperm-like objects (Fig. 4E) were shown to be nucleated when stained for DNA with DAPI (Fig. 4F), and were strikingly small in size when observed using Differential Interference Contrast (DIC) microscopy (∼ 1.5 μm diameter). To confirm that these small cells were indeed sperm, we applied an antibody that recognizes <u>M</u>ajor <u>S</u>perm <u>P</u>rotein (MSP). MSPs are small, highly conserved proteins in nematode sperm and are thought to have cytoskeletal and signaling functions (Roberts and Stewart, 1995; Miller *et al*., 2001). The monoclonal antibody was generated against the last 21 amino acids of the *C. elegans* protein MSP-40 (Miller *et al*., 2001) and has been successfully applied to *Acrobeloides* nematodes (Heger *et al*., 2010) which are in the same clade (IV) as *Steinernema* (Schiffer *et al*., 2018). To confirm that *msp* is conserved in *Steinernema spp,* we used WormBase ParaSite (Bolt *et al*., 2018) to search for *C. elegans msp-40* homologs in *S. carpocapsae,* the *Steinernema* species with the most complete genome (Dillman *et al*., 2015; Rougon-Cardoso *et al*., 2016; Serra *et al*., 2019). We found seven *msp* homologs in *S. carpocapsae* with 68.7-91.3% overall amino acid sequence identity to *C. elegans* MSP-40. Most importantly, the C-terminal 21 amino acids used to generate the monoclonal antibody we had used (REWFQGDGMVRRKNLPIEYNP) were 100% identical. We therefore expected that the sperm of *S. hermaphroditum* would contain this highly conserved antibody epitope and would be identified by antibody staining. Indeed, the small cells in the gonads of unmated animals with a female somatic appearance were confirmed by anti-MSP antibody staining to be hermaphroditic sperm. Thus, the structure they reside in is a spermatheca and the animal is hermaphroditic (Fig. 4D, G, H).

Altogether, we conclude that *S. hermaphroditum* is consistently hermaphroditic. Each generation consists almost exclusively of self-reproducing hermaphrodites, accompanied by spontaneous males. The mode of reproduction could be either self-fertilization (similar to that of *C. elegans)* or parthenogenesis.

### Forward genetic screens in *S. hermaphroditum* result in mutants with a wide variety of phenotypes

The consistent hermaphroditism of selected nematodes, in particular *C. elegans,* greatly facilitated the development of genetic tools (Brenner, 1974). Our discovery of the first consistently hermaphroditic entomopathogenic nematode motivated us to immediately develop forward-genetic tools in *S. hermaphroditum*. We adapted and optimized EMS mutagenesis protocols used on *C. elegans* to screen for mutant lines of *S. hermaphroditum*. A series of trial screens produced over three hundred candidate mutant animals resulting in at least thirty-seven independent mutant lines with visible and stable phenotypes with mostly 100% penetrance. The mutants showed a wide variety of phenotypes including dumpy (Dpy), various types of uncoordinated locomotion (Unc), gonad migration defective (Mig), dauer-constitutive (Daf), and high-incidence-of-males (Him) (Fig. 5, Table 1). Mutant strains were tested by mating with wild-type males to determine whether their phenotype was dominant or recessive; three strains could not be tested because of a highly variable phenotype or because they could not be successfully mated. Nearly all mutant phenotypes tested were recessive; the one exception, *unc(sy1654)*, had a recessive strong locomotion defect and a dominant phenotype that caused it to continuously twitch, strongly resembling loss-of-function mutants of the *C. elegans* gene *unc-22* (Moerman and Baillie, 1979).

**Table 1:** Description of mutant strains and alleles. A list of mutant *S. hermaphroditum* strains recovered in this work. Each mutant was recovered from different mutagenized P_0_ animals than any other mutant with the same phenotype; this includes one set of three mutants and another set of two mutants that fail to complement each other. Linkage group assignment was determined as described in Materials and Methods and as shown in Figure 5.

| Strain | Allele | Phenotype | Linkage Group | Description |
| --- | --- | --- | --- | --- |
| PS9258 | *mig(sy1637)* | Gonad migration defective | I |  |
| PS9255 | *dpy(sy1634)* | Dumpy, Vulvaless | Unknown | Unable to mate |
| PS9256 | *unc(sy1635)* | Uncoordinated | X | Reverse kinker |
| PS9257 | *unc(sy1636)* | Uncoordinated | III | Kinker; Fails to complement *unc(sy1647)* |
| PS9260 | *dpy(sy1639)* | Dumpy | X | Fails to complement *dpy(sy1644)* and *dpy(sy1646)* |
| PS9265 | *dpy(sy1644)* | Dumpy | X | Fails to complement *dpy(sy1639)* and *dpy(sy1646)* |
| PS9267 | *dpy(sy1646)* | Dumpy | X | Fails to complement *dpy(sy1639)* and *dpy(sy1644)* |
| PS9268 | *unc(sy1647)* | Uncoordinated | III | Kinker; fails to complement *unc(sy1636)* |
| PS9269 | *daf(sy1648)* | Dauer-constitutive | II |  |
| PS9270 | *unc(sy1649)* | Uncoordinated | Autosomal | Kinker. Slow growth. |
| PS9272 | *unc(sy1651)* | Uncoordinated | Autosomal | Kinker |
| PS9273 | *dpy(sy1652)* | Dumpy | III |  |
| PS9274 | *unc(sy1653)* | Uncoordinated | X | Kinker |
| PS9275 | *unc(sy1654)* | Uncoordinated | I | Dominant twitcher, recessive lethargic |
| PS9276 | *unc(sy1655)* | Uncoordinated | III | Reverse kinker |
| PS9277 | *dpy(sy1656)* | Small body size | Autosomal |  |
| PS9278 | *unc(sy1657)* | Uncoordinated | X | Reverse kinker |
| PS9279 | *dpy(sy1659)* | Dumpy | Autosomal |  |
| PS9282 | *unc(sy1661)* | Uncoordinated | Autosomal | Loopy |
| PS9283 | *dpy(sy1662)* | Dumpy | X | Complements *dpy(sy1639)*, *dpy(sy1644)*, and *dpy(sy1646)* |
| PS9285 | *dpy(sy1664)* | Dumpy | I |  |
| PS9287 | *dpy(sy1666)* | Dumpy | Unknown | No cross progeny from mating |
| PS9288 | *unc(sy1667)* | Uncoordinated | I | Lethargic, rigid posture |
| PS9289 | *unc(sy1668)* | Uncoordinated | II | Lethargic, rigid posture |
| PS9290 | *unc(sy1669)* | Uncoordinated | Autosomal | Kinker |
| PS9291 | *dpy(sy1670)* | Dumpy | Autosomal |  |
| PS9293 | *unc(sy1672)* | Uncoordinated | I | Reverse kinker |
| PS9294 | *mig(sy1673)* | Gonad migration defective | III |  |
| PS9295 | *unc(sy1674)* | Uncoordinated | Autosomal | Kinker |
| PS9296 | *dpy(sy1675)* | Dumpy | II |  |
| PS9300 | *unc(sy1679)* | Uncoordinated | Unknown | Loopy, variable phenotype |
| PS9301 | *him(sy1680)* | High incidence of males | Autosomal | Variable sickness, low brood size |

The progeny of crosses between wild-type males and mutant hermaphrodites were predominantly self-progeny rather than cross progeny, indicating that unlike what is observed in some other hermaphroditic nematodes such as *C. elegans* (Ward and Carrel, 1979) sperm introduced by mating with a male do not preferentially produce cross progeny in mated animals.

### *S. hermaphroditum* is androdioecious with chromosomal sex determination

Crosses of mutant hermaphrodites with wild-type males also revealed that sex determination in *S. hermaphroditum* is chromosomal: for 7 of 27 mutants, mating with wild-type males produced wild-type hermaphrodite cross progeny and mutant male cross progeny, in approximately equal numbers (47 hermaphrodites: 41 males and 116 hermaphrodites: 115 males for two independently recovered X-linked *dpy* mutants).

When double-mutant hermaphrodites homozygous for the autosomal mutation *him(sy1680)* and the X-linked recessive marker *unc(sy1635) X* were mated with wild-type males, phenotypically wild-type (non-Unc) male progeny were observed. A similar mating of wild-type males with *unc(sy1635) X* hermaphrodites produced cross-progeny that were all *unc(sy1635) X*/+ wild-type hermaphrodites and *unc(sy1635) X*/0 males; the observed non-Unc males from the cross with *him(sy1680)*; *unc(sy1635) X* hermaphrodites could only be explained by their containing a paternal wild-type X chromosome and not containing an *unc(sy1635)*-marked maternal X chromosome. This observation indicates that the high-incidence-of-males phenotype of *sy1680* results from X chromosome loss in oogenesis, further demonstrating the chromosomal nature of sex determination in *S. hermaphroditum*.

Although *S. hermaphroditum* animals consistently produce progeny without being mated, this is not enough to prove that it is a self-fertilizing hermaphrodite; an alternative explanation is that they could be producing self progeny through parthenogenesis. One way to test this is by examining the progeny of a heterozygous animal: if multiple markers show Mendelian ratios, this excludes the parthenogenesis model (See Fig. 6). To further examine if *S. hermaphroditum* is parthenogenic or self-fertilizing, we examined the self-progeny of animals heterozygous for each of several mutants with strong, easily detected mutant phenotypes. The F_2_ progeny of all mutants so examined segregated according to a Mendelian ratio (Table 2). In particular, two mutations, *dpy(sy1639)* and *unc(sy1635)*, that were both on the X chromosome and that were not closely linked to each other (approximately 15-20 map units; data not shown), both showed Mendelian segregation. These results are consistent with hermaphroditism by self-fertilization, and are not consistent with parthenogenesis.

**Figure 6:**
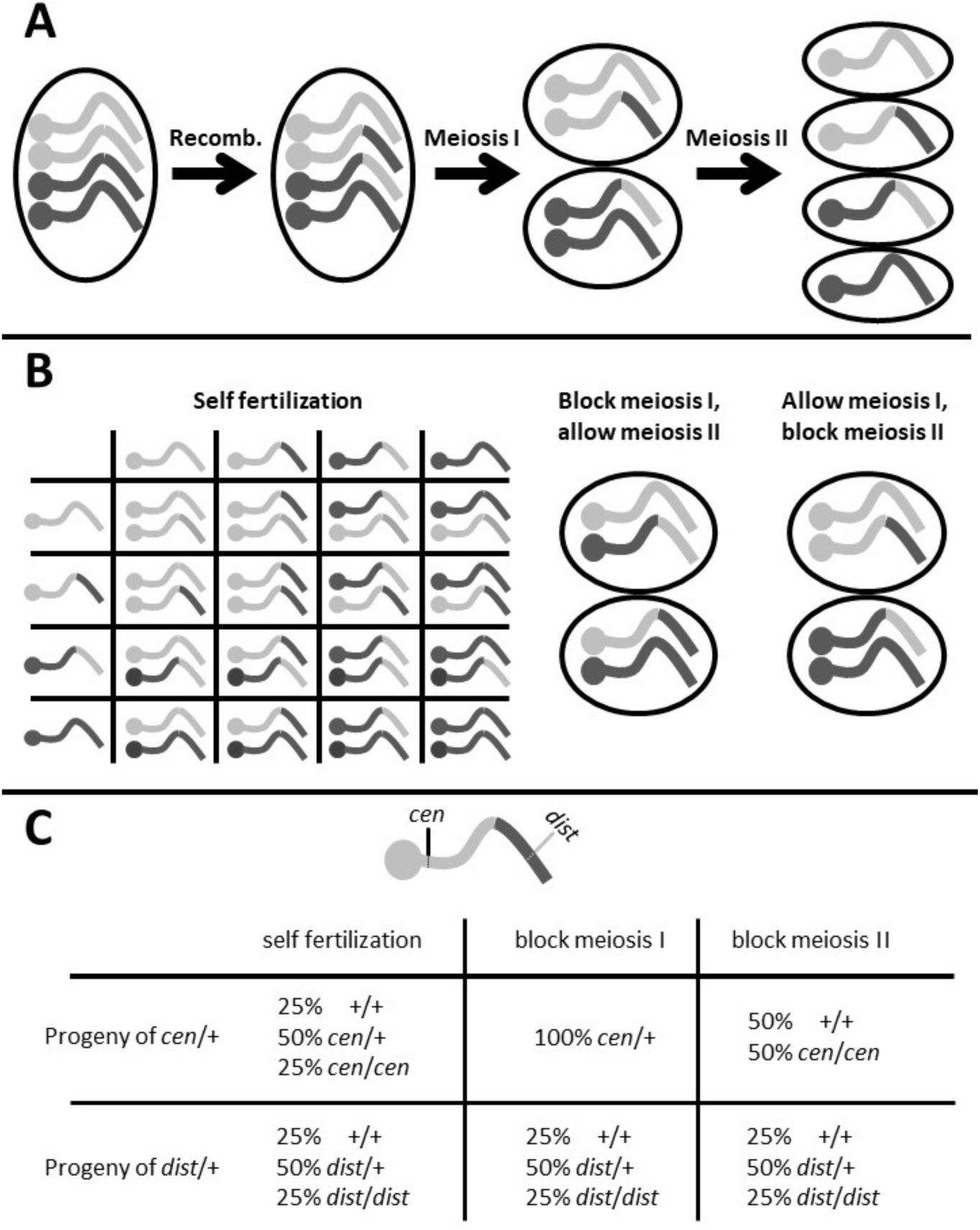
Parthenogenesis versus self-fertilizing hermaphroditism. Animals that have a grossly female appearance and that can reproduce without being mated might be producing progeny by self-fertilization or might be parthenogenic. The differences between these modes of reproduction would affect the suitability of the organism for genetic studies. (A) Normal meiosis is shown diagrammatically for a single chromosome, with its centromere visible in the cartoon as a ball at the left end, of the chromosome. This diploid animal has a pair of homologous chromosomes; one is shown as light in color, and one is shown as dark, representing their different haplotypes. Mitotic DNA replication has doubled the number of chromosomes during oogenesis. Each homologous chromosome is present in two copies of sister chromatids forming a tetraploid bivalent chromosome pair similar to Fig. 4I. Recombination can occur at this stage to cause the exchange of sections of the chromosome sequences distal to the site of crossing over with respect to the centromere, as shown for one pair in this cartoon. In Meiosis I the two homologous chromosomes are separated from each other, and in Meiosis II the individual sister chromatids are separated from each other. (B) In self-fertilization, a diploid embryo is generated by fusion of two haploid gametes produced by separate meiosis in oogenesis and spermatogenesis. In parthognogenesis, no sperm is engaged. The diploid embryo is produced by preventing the reduction of chromosome number in meiosis I or meiosis II in the oocytes. Examples are shown of the progeny classes that could result from this; focus on the different sorting of the centromeres into zygotes. (C) If we consider two markers, one tightly linked to the centromere (*cen*) and one (*dist*) distant from the centromere and genetically unlinked to it, and so recombinant with respect to the centromere half of the time. We can then examine the resulting progeny frequencies assuming the mechanisms shown in B: for the distal marker *dist* that isn’t genetically linked to the centromere no difference will be observed between mechanisms, but for the centromere-linked marker *cen* there will be a marked difference: 25% of self-fertilized embryos will be homozygous, but depending on the mechanism of parthenogenesis either 0% or 50% of progeny will be homozygous close to the centromere.

**Table 2:** Segregation of selected genetic markers from heterozygotes in self-reproduction. Animals heterozygous for various recessive markers were selfed and their progeny were sampled to determine what proportion were homozygous for the marker. In every case, that proportion was close to one in four. This is the result to be expected from self-fertilization; it is not consistent with parthenogenesis (see Figure 6). Note in particular that two of these markers, *unc(sy1635)* and *dpy(sy1639)* are approximately 20 map units apart on the same chromosome, suggesting that they cannot both be far distant from the centromere, as would be required to explain this proportion in parthenogenic reproduction.

| Marker | Proportion of self-progeny phenotypic (%) |
| --- | --- |
| *dpy(sy1664)* | 19/101 (19%)<br>33/139 (24%)<br>30/112 (27%) |
| *unc(sy1647)* | 26/107 (24%)<br>14/69 (20%)<br>29/102 (28%)<br>33/155 (21%)<br>37/148 (25%)<br>33/125 (26%) |
| *dpy(sy1652)* | 23/90 (26%)<br>28/108 (26%) |
| *dpy(sy1639)* | 38/136 (28%)<br>38/159 (24%)<br>29/123 (24%)<br>30/117 (26%) |
| *dpy(sy1644)* | 25/90 (28%)<br>12/68 (18%)<br>29/132 (22%) |
| *dpy(sy1646)* | 62/267 (23%)<br>25/106 (24%) |
| *unc(sy1635)* | 17/61 (28%)<br>30/120 (25%) |

### Identification of linkage groups in *S. hermaphroditum*

DAPI staining of gonads showed five pairs of tetraploid chromosomes in diakinesis stage oocytes (Fig. 3E), which is consistent with the chromosome numbers reported for other *Steinernema* species (Curran, 1989). We have begun using our collection of *S. hermaphroditum* mutants to generate the first genetic map of an entomopathogenic nematode (Fig. 7, Table 1). Genetic mapping has shown that mutations on the X chromosome are linked to each other (data not shown) and has thus far identified three autosomal linkage groups; mapping additional markers is expected to generate a map that matches our cytological observations.

**Figure 7:**
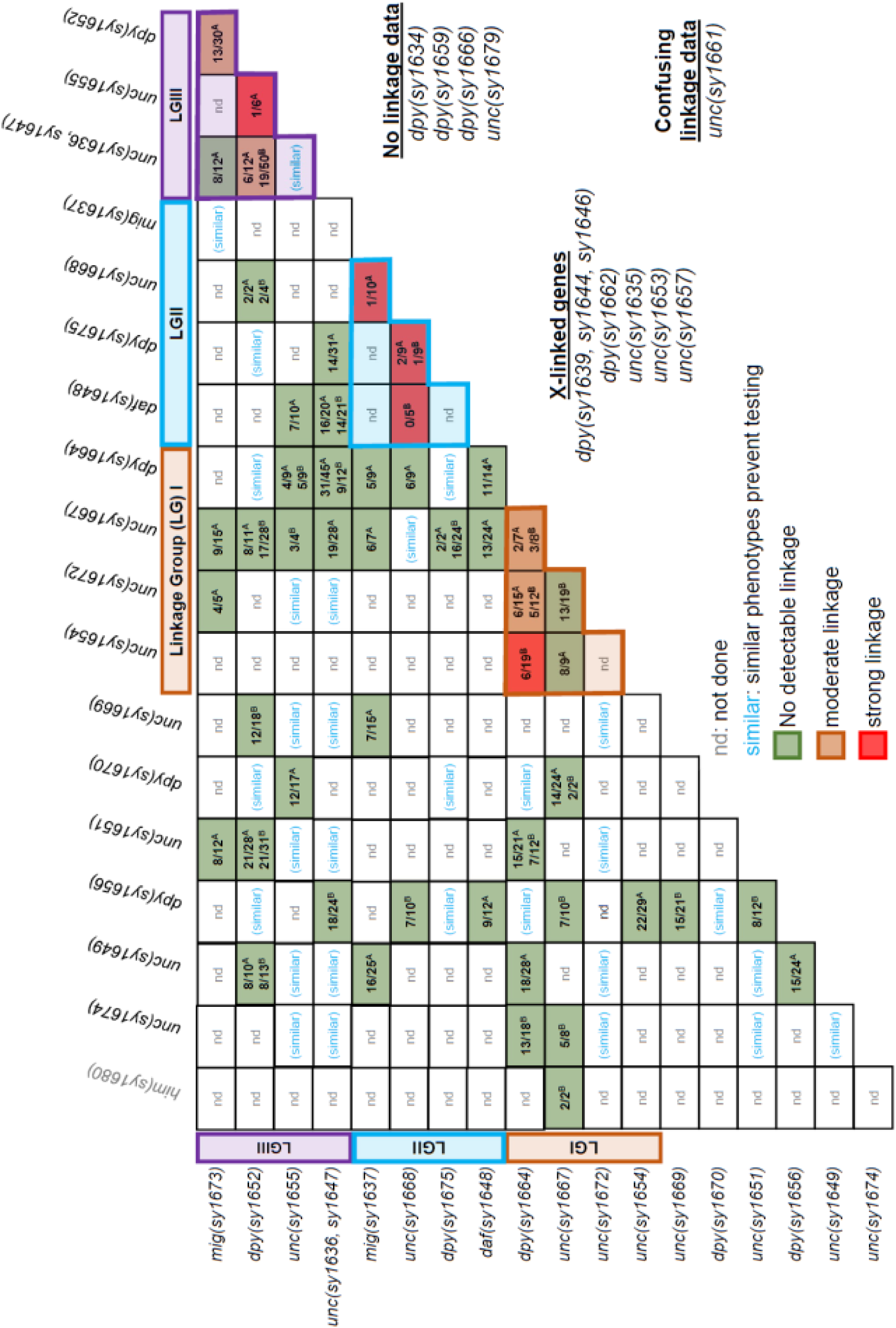
A first genetic linkage map of *S. hermaphroditum.* Genetic markers were mapped as described in Materials and Methods. Numbers shown are the proportion of homozygotes for a first marker whose progeny included animals showing the phenotype of a second marker; superscript “A” indicates the first marker was the mutation listed at left, labeling the row; superscript “B” indicates the first marker was the mutation listed above, labeling the column. Some mutations were difficult to score reliably, giving confusing results (*unc(sy1661)*, not included in the mapping data shown) or could be difficult to detect, giving the false impression of linkage (*dpy(sy1675)*, specifically in combination with *unc(sy1636)*). Seven mutations were assigned to the X chromosome, and were linked to each other on that chromosome (numbers not shown); 20 autosomal mutations have thus far defined three linkage groups, of an expected four. LG, Linkage Group.

## Discussion

In this research we report that a recently isolated Indian strain of *S. hermaphroditum* CS34 is a consistently hermaphroditic species. We have produced a highly inbred line of *S. hermaphroditum* to support molecular analysis and provide a consistent phenotypic basis for experimentation. The capacity for parasitism is retained in the inbred line, which remains able to efficiently infect and kill insect larvae in partnership with its *Xenorhabdis griffiniae* bacterial symbiont. We confirmed that hermaphrodites of this strain contain sperm and reproduce by self-fertilization, and can also be mated by males. EMS mutagenesis of this strain generated mutant lines that consistently demonstrate a variety of phenotypes and serve as genetic markers to demonstrate chromosomal sex determination and create an initial genetic linkage map of *S. hermaphroditum*.

Previous research characterized *S. hermaphroditum* as gonochoristic (male-female) after the first generation of growth in insects, based on an observation that animals in the second generation lacked sperm in their spermathecae (Stock *et al*., 2004). Through dissection and antibody staining, we identified sperm in the gonads of unmated animals with a female somatic morphology that had been cultured *in vitro* for over 10 generations. Previous microscopic examination done without the assistance of an antibody may have overlooked these sperms due to their small appearance using DIC (LaMunyon and Ward, 1998). Alternatively, the discrepancy between our findings and the previous report might reflect differing environmental factors, such as the effects of growing multiple generations *in vitro* on cultured bacteria, rather than infecting insects. In addition, the determination that the second generation of nematodes after infecting an insect consisted of females rather than hermaphrodites was from examination of a different *S. hermaphroditum* isolate, one collected more than 15 years earlier and nearly 10,000 kilometers away (Stock *et al*., 2004); the two strains may have differed in this regard. These questions could be addressed by studying a collection of more diverse wild isolates to explore the natural diversity within *S. hermaphroditum*.

Traditional EPN *in vitro* growth methods using lipid agar and liver-kidney agar have been widely applied in nematology and microbial symbiosis research. These rich media could support nematode growth either with symbiotic bacteria or, in the latter case, without bacteria (axenically) (Surrey and Davies, 1996; Vivas and Goodrich-Blair, 2001; Flores-Lara *et al*., 2007) and facilitated the description of the animals and the mass production of nematodes, especially IJs. However, they also hampered the application of basic techniques required for molecular and experimental genetics. The presence of animal tissue in the media decreases visibility of the nematodes, while oil droplets hamper nematode picking and cause inconsistency through variable distribution in the media. We adapted standard growth conditions (NGM) for laboratory culture of the free-living soil nematode *C. elegans* to grow both *S. hermaphroditum* and its *Xenorhabdus* bacterial symbiont and demonstrated that basic techniques developed in *C. elegans* could be adapted to this entomopathogenic nematode. Our purpose in adopting *C. elegans* techniques to encourage the application of approaches from multiple research traditions to better serve various functions in a broader research community.

*Xenorhabdus* bacteria undergo phenotypic switching phenomena: pleiotropic variation of observable phenotypes such as secondary metabolite production and motility (1° to 2° form) (Boemare and Akurst, 1988) and virulence modulation, in which individual cells can become attenuated for virulence in insects (Park *et al*., 2007), each of which can affect interactions between bacteria and nematodes. *S. carpocapsae* newly isolated from the wild are reported to associate exclusively with the 1° form of their *X. nematophila* symbiont (Forst *et al*., 1997). In virulence modulation, virulence-attenuated-phase bacteria colonize their nematode host and support its reproduction better than do the same bacterial species in a more virulent phenotypic phase (Cao *et al*., 2017). In this research, we isolated ten colonies of symbiotic bacteria *X. griffiniae* from a natural population of *S. hermaphroditum* IJs and found the production of brown pigment varied among the isolates. Pigmented bacteria supported nematode reproduction better than non-pigmented isolates did, suggesting potentially important consequences of phenotypic switching in this species. The various phenotypes of individual isolates of symbiotic bacteria that have correspondingly different impacts on the physiology of their host nematodes provoke intriguing questions about the possible heterogeneity of both nematode and bacteria in natural populations. Multiple closely-related strains of *X. griffiniae*, not distinguishable in the limited sequencing we performed may be associated with a natural population of nematodes; we have not yet observed spontaneous changes in bacterial phenotype in the isolates we have worked with.

The ability of *Xenorhabdus* bacteria to support the reproduction of their native host *Steinernema* nematodes is a crucial aspect of their mutualistic relationship (Herbert and Goodrich-Blair, 2007). We monitored *S. hermaphroditum* growth using as its food source either its native symbiont or three bacterial strains commonly used in *C. elegans* research. The ability of *E. coli* (OP50 and HB101) to support the development of *C. elegans* but not to robustly support *S. hermaphroditum* growth indicates these two bacteriovorous nematode species may have different nutritional needs or food preferences as a result of their distinct life styles (free-living vs parasitic). Unexpectedly, we discovered that a non-symbiotic bacterial species *Comamonas aq* could support nematode growth better than its native symbiotic bacterium *X. griffiniae.* Therefore *Comamonas aq* could serve as a neutral (non-pathogenic and non-mutualistic) food source for *S. hermaphroditum.* However, the advantage of *Comamonas* in supporting *S. hermaphroditum* growth *in vitro* does not necessarily mean *Comamonas* would be a better partner in the wild. Firstly, the *in vitro* growth conditions may not represent those *in vivo*. The insect carcasses are normally abundant sources of selected nutrients and the entomopathogenic nematodes would therefore have less need for their bacteria to abundantly produce those nutrients; also, the insect carcass might better enable the *X. griffiniae* metabolism to support nematode growth than does the agar media used in our assays. Even more importantly, *Xenorhabdus* bacteria contribute to insect-killing and help protect the carcass from other predators and other microbial species by producing inhibitory molecules, such as toxins and secondary metabolites with antimicrobial effects (Herbert and Goodrich-Blair, 2007). The antagonizing mechanisms protect the bacterial and nematode partners from exposure to potentially invasive microbes and ensure the fitness and fidelity of their mutualistic symbiosis (Herbert and Goodrich-Blair, 2007). These antagonizing mechanisms of the bacteria may also moderately inhibit nematode growth. Future research can take advantage of the unique binary symbiosis system of *S. hermaphroditum* and *X. griffiniae* to investigate the role of native symbionts and other environmental microbes at different stages of the symbiotic life cycle, such as nematode-bacteria signaling during the infective stage. As *S. hermaphroditum* is established into a genetic model it will be possible to study the mechanisms that tie these two symbiotic partners together, experimentally modifying both partners in the interaction.

We found EMS mutagenesis to be an efficient way of recovering mutant *S. hermaphroditum* strains with highly specific defects in diverse aspects of their biology. Mutants could easily be maintained and could be used to demonstrate a number of basic but essential features of this species. We found that sex is determined chromosomally, by the presence or absence of a second X chromosome. Sex determination appears to be XX/XO as opposed to XX/XY, because hermaphrodites can spontaneously generate male progeny. In addition, we showed that a Him mutant greatly increased the frequency of spontaneous male progeny by generating oocytes that lacked an X chromosome. By mating of wild-type males to mutants with recessive phenotypes that allowed the identification of cross-progeny we demonstrated that *S. hermaphroditum* does not exhibit a strong preference for the use of male-derived sperm. This phenomenon has been a notable feature of *C. elegans* biology and has been proposed to promote the small amount of outcrossing necessary to prevent uniformity in an inbred population (Ward and Carrel, 1979). Critical for future genetic research using *S. hermaphroditum*, we could use visibly phenotypic mutants to show that self-progeny are produced by fertilization with sperm that the hermaphrodites produce, rather than by parthenogenesis. In parthenogenesis, the cell that would normally develop into a haploid oocyte instead becomes a diploid zygote, without being fertilized by sperm. Different mechanisms can achieve this result, including preventing the reduction of chromosome number at meiosis I or meiosis II; any such mechanism will result in different frequencies of homozygotes segregating from a mother heterozygous for a marker, depending on the genetic distance between that marker and the chromosome’s centromere (Fig. 6). We found that for every marker tested, including two markers distant from each other on the same chromosome, homozygotes were one quarter of the progeny – a Mendelian ratio that is the hallmark of self-fertilization. This ensures that it will be possible to screen efficiently for recessive phenotypes at all points in the genome.

The collection of mutants we have already assembled is sufficient to describe how sex is determined in *S. hermaphroditum* and offers a proof of principle of the potential of forward genetic screens in this organism. Future screens will be able to target biological questions involving the unique biology of the entomopathogenic nematode, such as the recognition, maintenance, and transmission of its mutualistic symbiont and its ability to hunt and to parasitize insects. Already we have a couple of interesting mutants: one that in every generation causes its progeny to develop as infective juveniles rather than prioritize reproduction, and two unlinked mutations sharing a novel phenotype of altered gonad shape that may reveal fundamental mechanisms of cell migration. Development of transgenesis and of reverse-genetics tools such as RNAi and CRISPR-Cas9 genome editing should expand a genetics toolkit we have begun by exploring gene function through forward genetics. Engineered nucleases such as CRISPR-Cas9 can also be used to build the tools that will make forward genetics more powerful (Dejima *et al*., 2018). The development of a complete and annotated *S. hermaphroditum* genome would greatly facilitate both the targeting of genes for reverse genetics and the identification of genes mutated in forward genetic studies. We hope that the discovery of additional wild isolates of *S. hermaphroditum* will facilitate genetic mapping in this species, enable the study of this species’s natural diversity, potentially including molecular pathways underlying sex determination, and comparative study among free-living and parasitic nematodes.

### Data and reagent availability

Strains described in this work are available upon request. 16S rRNA gene sequences of the *X. griffiniae* bacterial symbionts of the *S. hermaphroditum* isolate CS34 are available in the Genbank database and the accession numbers for 16S rRNA sequence (including HGB2511) are MZ913116-MZ913125.

## Acknowledgments

We are overwhelmingly grateful for *S. hermaphroditum* as a generous gift from nature and a superb organism to explore scientific curiosities. We thank Adler Dillman of University of California Riverside for generously sharing the *S. hermaphroditum-*India strain CS34 and for comments on this manuscript. We also thank Heidi Goodrich-Blair and Jennifer Heppert of the University of Tennessee Knoxville for helpful discussions and editorial suggestions. We are also grateful for valuable suggestions from Zuzana Kocsisova. The monoclonal anti-MSP antibody 4A5 developed by David Greenstein of the University of Minnesota was obtained from the Developmental Studies Hybridoma Bank, created by the NICHD of the NIH and maintained at The University of Iowa Department of Biology. Bacterial strains HB101 and DA1877 were obtained from the *Caenorhabditis* Genetics Center (CGC), which is funded by the NIH Office of Research Infrastructure Programs (P40 OD010440). This work was also facilitated by WormBase and ParaSite, a knowledgebase for nematode research. This research was supported by NIH-NRSA F32 5F32GM131570-03 (MC); NSF EDGE grant 2128267, and the Center for Evolutionary Science at Caltech.

## Supplemental Data

**Fig. S1.**
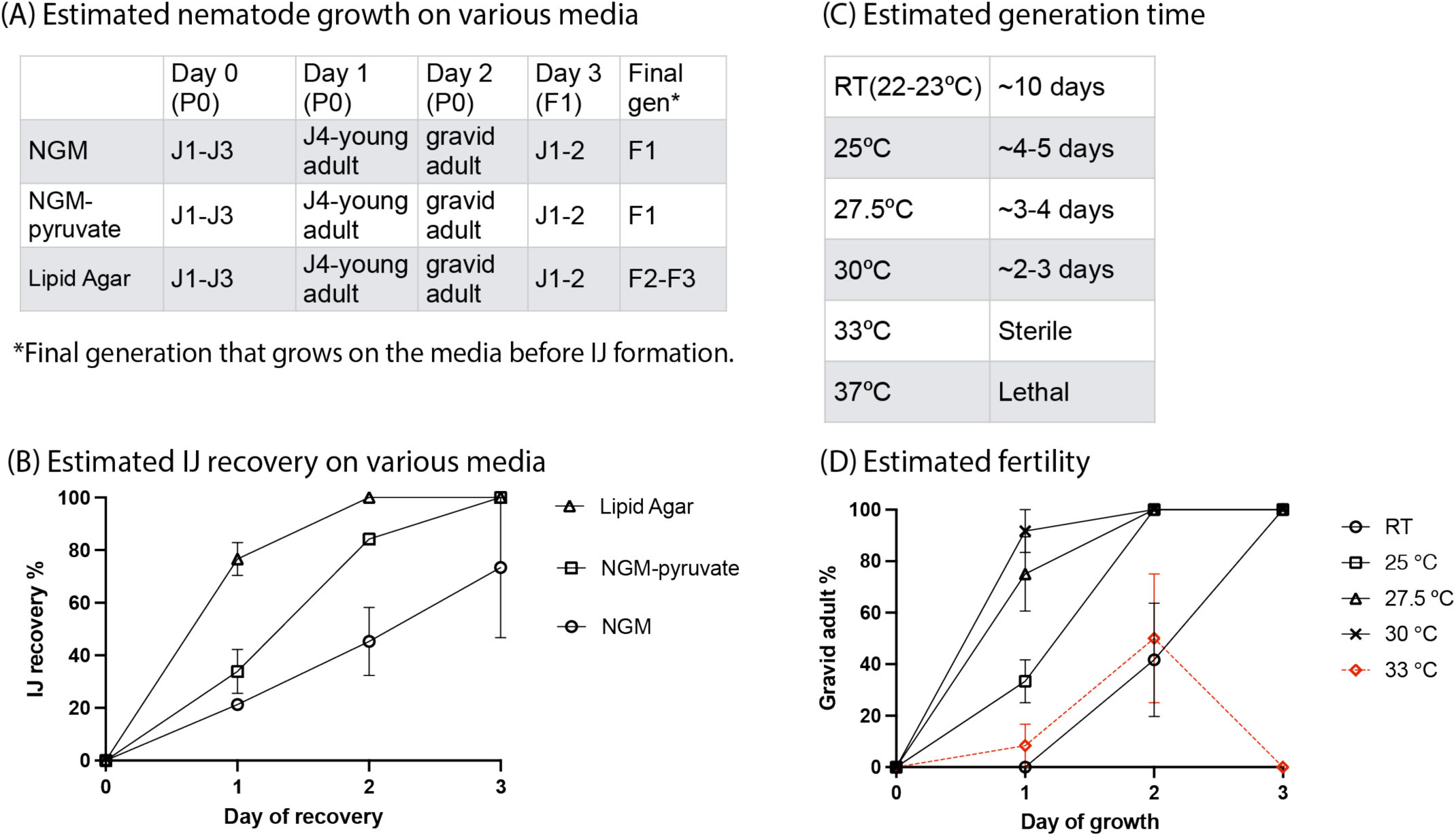
**The effect of media choice and temperature on *S. hermaphroditum* growth.** (A): Approximate nematode life cycle progression on NGM, NGM supplemented with pyruvate, and lipid agar supplemented with cholesterol at 25°C. A small population of young juveniles (J1-J3) were transferred to lawns of their bacterial symbiote HGB 2511 on Petri plates and estimated for life stage progression daily. The representative observations from three replicates are shown (B): Estimated IJ recovery on NGM, NGM supplemented with pyruvate, and lipid agar. Approximately 10-20 individual IJs were transferred onto a Petri plate containing the indicated media with a lawn of HGB2511 and incubated at 25°C. IJ recovery was monitored daily and percent IJ recovery was calculated by recovered IJs out of the total number of recovered and non-recovered IJs. The averages and standard errors of three replicates are shown. (C). Estimated generation time at various temperatures. Animals were grown on lawns of their bacterial symbiont HGB2511 on NGM agar. Sterile indicates animals remained alive but failed to produce progeny; lethal indicates animals died early and without producing progeny. (D). Estimated fertility at various temperatures. Four young adults were transferred onto lawns of their bacterial symbiont HGB2511 grown on NGM agar and incubated at room temperature (RT), 20-22°C; 25°C, 27.5°C; or 33°C. Gravid adults were identified by the presence of embryos in the gonads and percentage of gravid adults in the total population was monitored and calculated daily as an indication of fertility. The averages and standard errors of three replicates are shown.

**Fig. S2:**
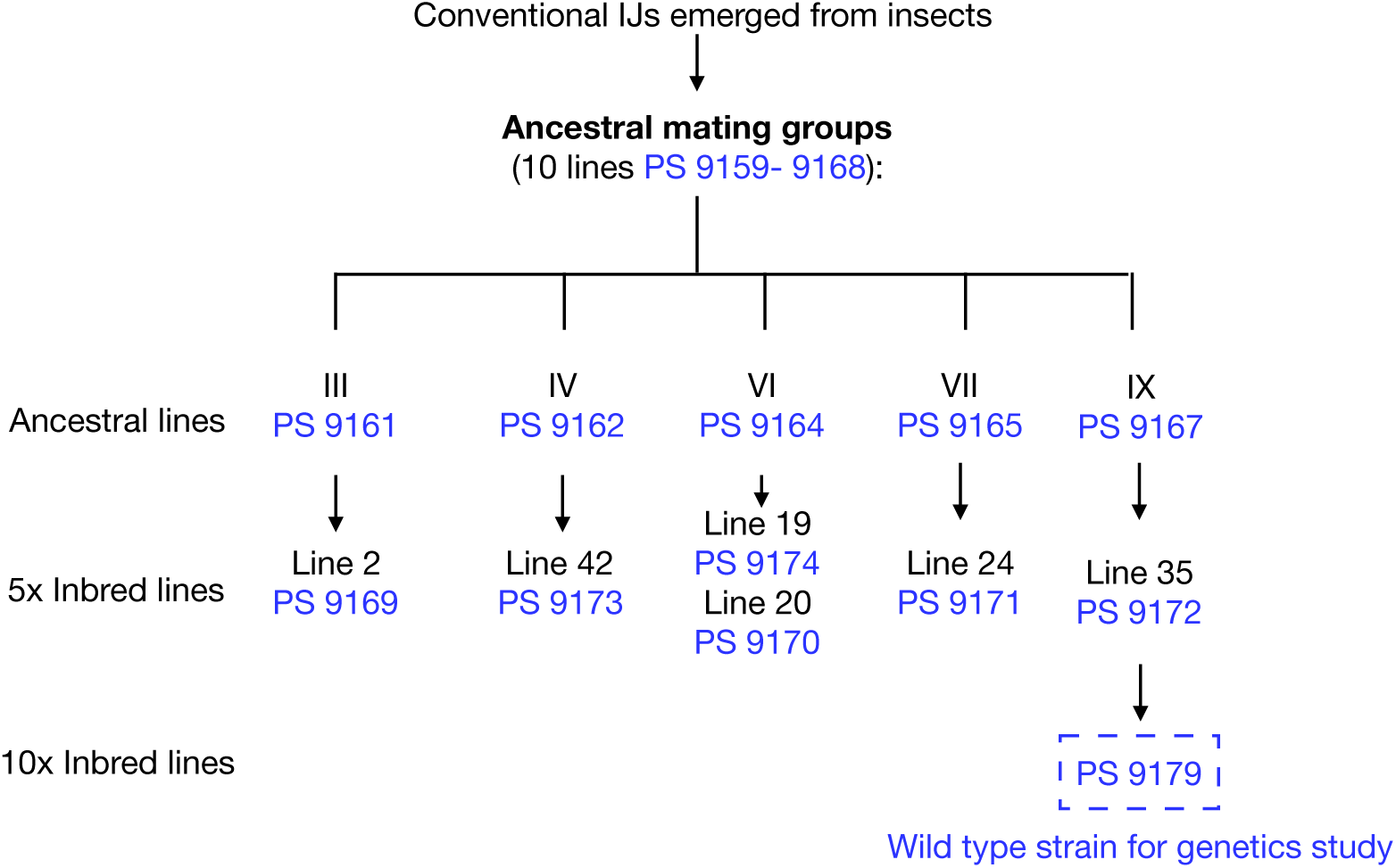
Inbreeding of *S. hermaphroditum* wild isolate CS34 to generate a wild-type reference strain with a stable phenotype and a homozygous genetic background. Conventional IJs recovered from infected insects were cultured *in vitro* (on lipid agar) to establish ten ancestral mating groups (Ancestral lines I-X). The ancestral line number is determined by the symbiont isolate number with which the line was co-cultured (Isolate 1-10, corresponding to the ancestral line with the same number, I to X). Five ancestral lines were chosen to further produce inbred lines (Ancestral group III, IV, VI, VII, and IX). In each generation, ten individual unmated hermaphrodites each were placed on a Petri plate containing NGM agar seeded with the corresponding bacterial isolate of *X. griffiniae*. The diagram shows the ten lines of ancestral mating groups, six lines inbred for five generations, and one line originated from Ancestral IX inbred for ten generations. The resulting line PS9179 was used for further study. It was grown on bacterial isolate 9, later renamed HGB2511.

**Table S1:** Bacteria and wild-type nematode strains used in this study*.

| Strain | Description | Source |
| --- | --- | --- |
| HGB2511 | *Xenorhabdus griffinae* native symbiont isolated from *S. hermaphroditum* CS34. | This study |
| OP50 | *Escherichia coli* uracil auxotroph and conventional food source for *C. elegans*. | Brenner 1974. |
| HB101 | *Escherichia coli* mutant and commonly used as a food source for *C. elegans*. | (Boyer and Roulland-dussoix, 1969) |
| DA1877 | *Comamonas aquatica*, a commonly used food source for *C. elegans*. | (Shtonda and Avery, 2006; Watson *et al.*, 2014) |
| CS34 | *S. hermaphroditum*-India wild isolate. | (Bhat *et al.*, 2019) |
| PS9167 | Ancestral group IX of CS34 inbred line co-cultured with *X. griffinae* (HGB 2511). | This study |
| PS9172 | Derived from Ancestral group IX (PS9167) and inbred for five generations. | This study |
| PS9179 | Derived from Ancestral group IX (PS9167) and inbred for ten generations. Used as wild type for the genetic study in this research. | This study |

\*See Table 1 for a list and description of the mutant strains produced in this study. See Figure S3 for a diagram of the inbreeding of *S. hermaphroditum* that resulted in the strain PS9179.

**Table S2:**
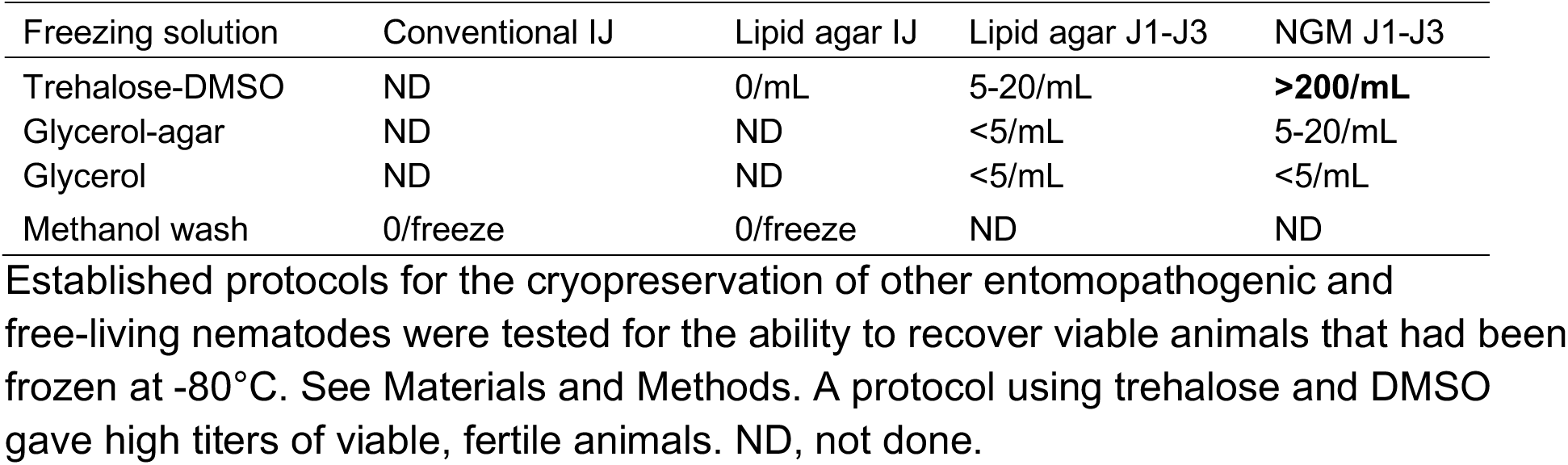
Cryopreservation of *S. hermaphroditum*.

**Movie: S1:** A movie showing the twitcher phenotype of the uncoordinated mutant *unc*(*sy1654*). https://caltech.box.com/s/c3j4v12r8bbzflwclsibfv8hoem8ldv0

**Movie: S2:** A movie showing the reverse kinker phenotype of the uncoordinated mutant *unc(sy1635)*. https://caltech.box.com/s/ec77sjeznct2bdwbhzq61nfbg6f99u3q

